# Benchmarking tools for transcription factor prioritization

**DOI:** 10.1101/2024.04.23.590206

**Authors:** Leonor Schubert Santana, Alejandro Reyes, Sebastian Hoersch, Enrico Ferrero, Christian Kolter, Swann Gaulis, Sebastian Steinhauser

## Abstract

Spatiotemporal regulation of gene expression is controlled by transcription factor (TF) binding to regulatory elements, resulting in a plethora of cell types and cell states from the same genetic information. Due to the importance of regulatory elements, various sequencing methods have been developed to localise them in genomes, for example using ChIP-seq profiling of the histone mark H3K27ac that marks active regulatory regions. Moreover, multiple tools have been developed to predict TF binding to these regulatory elements based on DNA sequence. As altered gene expression is a hallmark of disease phenotypes, identifying TFs driving such gene expression programs is critical for the identification of novel drug targets.

In this study, we curated 84 chromatin profiling experiments (H3K27ac ChIP-seq) where TFs were perturbed through e.g., genetic knockout or overexpression. We ran nine published tools to prioritize TFs using these real-world data sets and evaluated the performance of the methods in identifying the perturbed TFs. This allowed the nomination of three frontrunner tools, namely RcisTarget, MEIRLOP and monaLisa. Our analyses revealed opportunities and commonalities of tools that will help to guide further improvements and developments in the field.

## Introduction

Spatiotemporal gene expression levels are regulated by binding of transcription factors (TFs) to regulatory elements [1]. TF binding is regulated by various factors such as DNA accessibility, epigenetic factors (e.g., DNA methylation) and co-factor binding [2–4]. Further, TFs link cellular signalling pathways to gene expression programs which in turn regulate specific cellular actions (e.g., differentiation, apoptosis) [5]. Hence, gene regulation is fundamental for the plethora of cell types in complex organisms, and regulatory alterations are a common denominator for various diseases [6].

Several high-throughput sequencing methods have been developed to interrogate the different layers of transcriptional regulation including gene expression (e.g., RNA-seq) and regulatory elements (e.g., Assay for Transposase-Accessible Chromatin using sequencing (ATAC-seq) or chromatin immunoprecipitation followed by sequencing (ChIP-seq)) [1]. Genome-wide mapping of the acetylation of lysin 27 in the H3 histone (H3K27ac) is commonly used to identify active regulatory elements, such as enhancers and promoters [7]. Moreover, wide-spread enrichments of H3K27ac along large consecutive genomic locations have been used to define super-enhancers (SEs), which are postulated to be important regulators of cell identity genes [8]. However, it remains controversial if SE are different from other regulatory elements such as enhancer clusters or holo-enhancers [9,10].

Many studies have used H3K27ac to investigate differences in regulatory element activity between experimental conditions (e.g., healthy vs disease phenotype or control vs compound treatment) [11–15]. A common downstream analysis based on differential regulatory elements is the identification of TFs which bind to these elements and therefore might play an important role in the observed phenotypes. Usually, the top-raking TFs in such analyses are used to formulate hypotheses that are further validated experimentally (e.g., by RNAi knockdown, knockout, compound modulation).

To this end, computational tools have been developed to perform TF prioritization based on different assumptions and implementations [16–24]. Among these, we could broadly identify two types, depending on their underlying reference: 1) tools leveraging DNA sequence information using position weight matrices (PWMs) to predict TF binding (PWM based tools), and 2) sequence-independent tools using previously identified TF binding sites in the genome (ChIP-seq peak based tools). Independently of their reference, both types of tools are prioritizing TFs based on statistical methods such as Fisher’s exact test, rank based enrichment, and LASSO regression, among others [16–24].

Although these tools play an important role for hypothesis generation in the scientific community, to our knowledge they have not been benchmarked for their ability to prioritize TFs.

In this study, we set out to identify the TF prioritization tools that yield the most accurate results, thus helping to formulate hypotheses for experimental validation with a higher probability of success. For this purpose, we are introducing a benchmarking framework based on the combination of 84 published H3K27ac ChIP-seq data sets with nine different TF prioritization tools. All selected H3K27ac ChIP-seq data sets included at least one TF perturbation (e.g., overexpression (OE), knockdown (KD)) providing us with a ground truth for each data set (TF labels). We ran each tool on all selected data sets, converted the tool outputs into TF priority rankings, and examined the tool performance using these TF labels against eight performance metrics. Finally, we investigated the importance of experimental variables on tool performance using random forest classifiers to model the tool results.

In summary, we present a benchmark study of TF prioritization tools based on real world data sets and give recommendations about tool selection highlighting potential improvements for new ones.

## Results

### A benchmarking framework to access the performance of TF prioritization tools

We designed a benchmarking framework for TF prioritization tools based on 84 publicly available H3K27ac ChIP-seq experiments from 53 different studies (Fig. 1 and Table S1). These data sets were selected based on the following criteria: 1) the raw data were available in the Gene Expression Omnibus (GEO), 2) the H3K27ac ChIP-seq assay was performed in human or mouse samples, and 3) the experimental design included at least one TF perturbation with corresponding control condition.

**Fig. 1.**
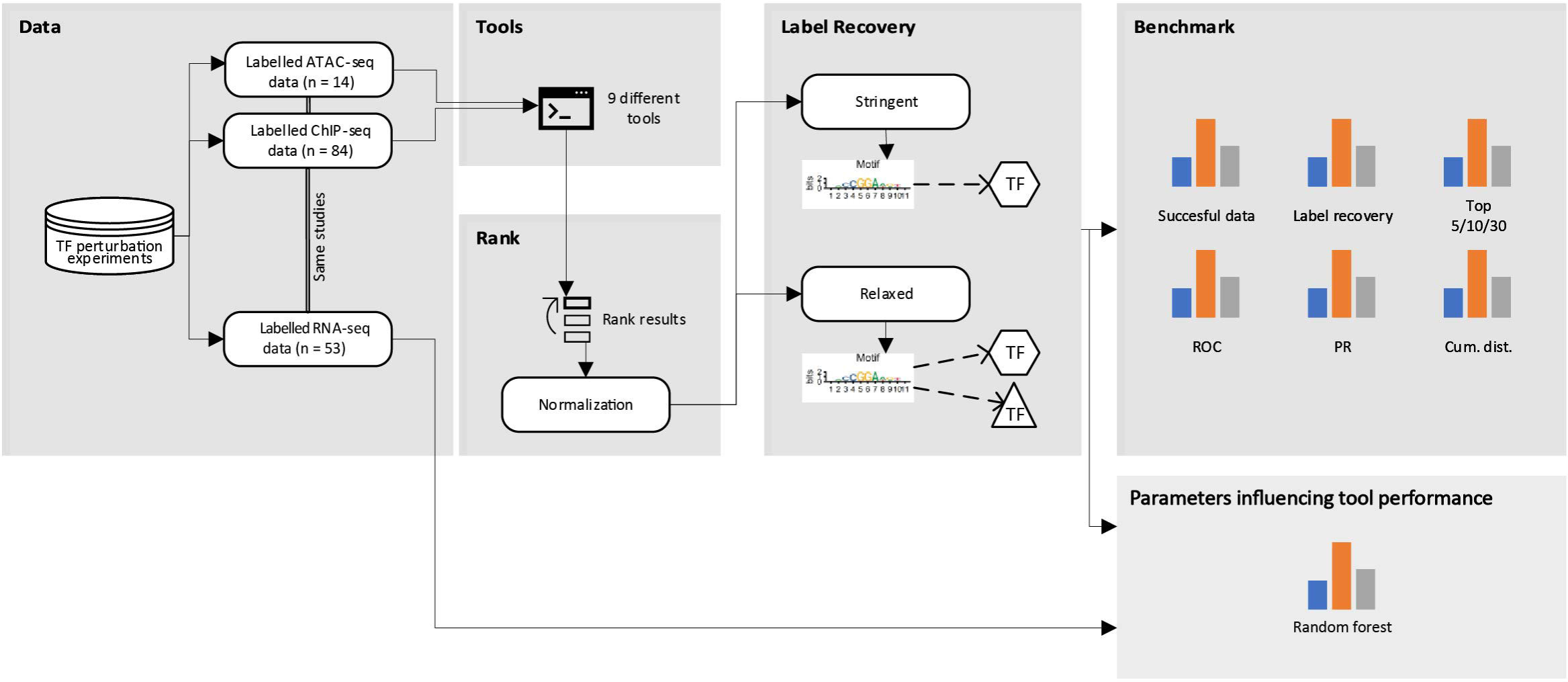
Schematic of the benchmarking framework to access the performance of TF prioritization tools. Data curation step: manual data set curation of H3K27ac experiments with underlying TF perturbation (e.g., TF knockout or over-expression), yielding 84 ChIP-seq data sets, a subset of 53 with matched RNA-seq and 13 with matched ATAC-seq. Tool implementation step: implementation of nine TF prioritization tools and inference of TFs on the 84 data sets. Ranking step: Resulting outputs are converted to ranked TF lists based on the tool statistic (e.g., p-value, AUC, or Z-score). Rankings are scaled to values between 0 and 1 (see Methods) to ensure cross tool comparability. Label recovery step: The scaled rankings are searched for the first occurrence of the experiment label (= perturbed TF). This analysis was performed using either a stringent label definition (exact TF match) or a more relaxed definition (any TF binding a similar motif). Benchmark step: These label recovery strategies in combination with the resulting rankings were used to compute eight benchmark metrics for each of the tools.

Using these criteria, we identified 40 mouse and 44 human experiments from tissues (n=17), primary cells (n=12), or immortalized cell lines (n=55), in which a TF was perturbed either by a knockout (KO, n=33), knockdown (KD, n=15), overexpression (OE, n=21) or compound treatment (either agonist or antagonist, n=15; Fig. S1A). Together, these experiments covered diverse characteristics reflecting standard experimental settings (Fig. S1). For example, the underlying ChIP-seq experiments were performed using three common commercially available H3K27ac antibodies and were sequenced from one to up to five replicates (Fig. S1B, C and Table S1).

Overall, our perturbed TF data sets cover 18 TF families out of the 66 defined by Lambert *et al.* (Fig. S1D) [2]. Notably, the most prominent TFs profiled were nuclear receptors (e.g., NR1H2, AR, PPARA), zinc finger TFs (e.g., KLF4, BCL6, EGR1) and GATA factors (TRPS1, GATA3, GATA4). The most common experimental design was the perturbation of nuclear receptors in mouse profiled with the Abcam ab4729 antibody (Fig. S1E).

We performed a literature search to identify candidate tools for TF binding prediction using the following inclusion criteria: 1) H3K27ac ChIP-seq data could be used as input, 2) the underlying code was available and useable either as command line tool, R, or Python package, 3) the code was published using a free and open-source licence. This led us to nine tools which can be categorized by basic principle into PWM- (n=7) and ChIP-seq peak-based (n=2). Moreover, the tools can be classified by the prioritization strategy into enrichment- (n=5), regression- (n=2), graph- (n=1) and ensemble-based (n=1; Fig. S2, Table S2) [16–24]. [16–24]In addition, some tools make specific biological assumptions; for example, CRCmapper is aiming to map core regulatory circuits (CRCs), which in turn are based on the existence of super-enhancers [8,20].

We applied all nine tools (where possible with multiple PWM libraries and backgrounds) to perform TF prioritization using the 84 H3K27ac ChIP-seq data sets as input. This resulted in 13 different TF prioritization approaches.

To compare the performance of the different approaches, we converted the metric of each approach (e.g., p-value, AUC, or z-score) into scaled ranks (Fig. 1). For tools outputting multiple ranking metrics, we chose the best performing metric for each of the tools (see Methods, Fig. S5A, B).

We examined the two most common parameters, the PWM motif library and the set of background sequences used by a tool. These parameters were only explored where accessible via command line arguments. For the PWM motif library, we compared the default motif libraries of a given tool with a recently published consensus library containing 5,594 PWMs covering 1,210 TFs (referred to as “+Lambert”) [2]. For the tools that enabled to change the background sets, we reported the tools background default and a background based on genomic regions where H3K27ac was enriched in the control conditions compared to the perturbed condition (referred to as “+bg”, for background). However, we made a comparative analysis of different backgrounds and found that the influence of the background set is neglectable compared to the TF tool, the ranking metrics of the different tools and the TF library (Fig. S5).

Throughout the manuscript, we refer to the perturbed TF in an experiment as the experiment label. We considered two criteria to assess whether the perturbed TF could be recovered from the data. For the *stringent* criterion, we required the TF *name* associated with a particular PWM/peak set to be the same as the TF perturbed in the experiment. For the *relaxed* criterion, we required the ranked PWMs/peak sets to be associated with a TF *homologous* to the perturbed TF (e.g., GATA1 PWM for GATA2 as label). The main rationale for the relaxed criterion was to allow for a fair comparison of approaches using different PWM/ChIP-seq peak collections and to address PWM redundancy between homologous TFs. The recovered TF labels in combination with the scaled rankings were used to compute eight different metrics (Fig. 1, Methods, and next Results section).

In summary, we assembled a diverse set of TF prioritization tools and combined with a representative set of TF perturbation H3K27ac ChIP-seq experiments into a benchmarking framework to examine their performance on real world experimental data.

### Benchmark comparison of TF prioritization tools based on recovering perturbed TFs

To exclude the possibility of systematic bias introduced by the data sets, we investigated the number of tools that returned the perturbed TF in the results. For clarity, we named a label recovered if the perturbed TF is shown at all in the output of a tool. We observed that for all 84 data sets, at least two tools returned the expected TF label (Fig. S3A, B). For 72 of them, the TF label ranked among the top 30 for at least one tool using the relaxed label recovery criterion (Fig. S3B).

Next, we benchmarked the TF prioritization tools using eight different metrics (see Methods). The first metric we computed for each tool was the number of data sets processed without errors and the number of TF labels recovered using both the stringent and relaxed criteria. Only four of the TF prioritization approaches did not complete for all 84 experiments using default parameters (Fig. 2A, B, white bars). TFEA, GimmeMotifs, HOMER + Lambert + bg and HOMER + bg failed to run for 26% (22), 21% (18), 7% (6) and 1% (1) of the experiments, respectively. Frontrunners using this metric were RcisTarget with and without background, which recovered 82 labels, and HOMER + Lambert, MEIRLOP and monaLisa with 81 labels recovered (Fig. 2B). The tools with the least recovered labels were CRCmapper (n=54) and LOLA (n=49). The stringent label recovery strategy gave a similar ranking on performance with fewer labels recovered overall (Fig. 2A, median number of TFs recovered, stringent n=73 and relaxed n=75). Only RcisTarget + bg (n=82) performed the same as in the relaxed strategy. The second-best approaches were HOMER + Lambert (n=80) and monaLisa (n=80), both recovering one fewer TF label than using the relaxed strategy. The tools with the lowest recovery were again CRCmapper and LOLA which only reported 37 and 42 labels in their results, respectively. Overall, none of the tools recovered all 84 TF labels and each label was recovered by at least two tools, suggesting that the label recovery failures were not driven by specific datasets but rather were tool-specific (Fig. 2A, B and Fig. S3A, B).

**Fig. 2.**
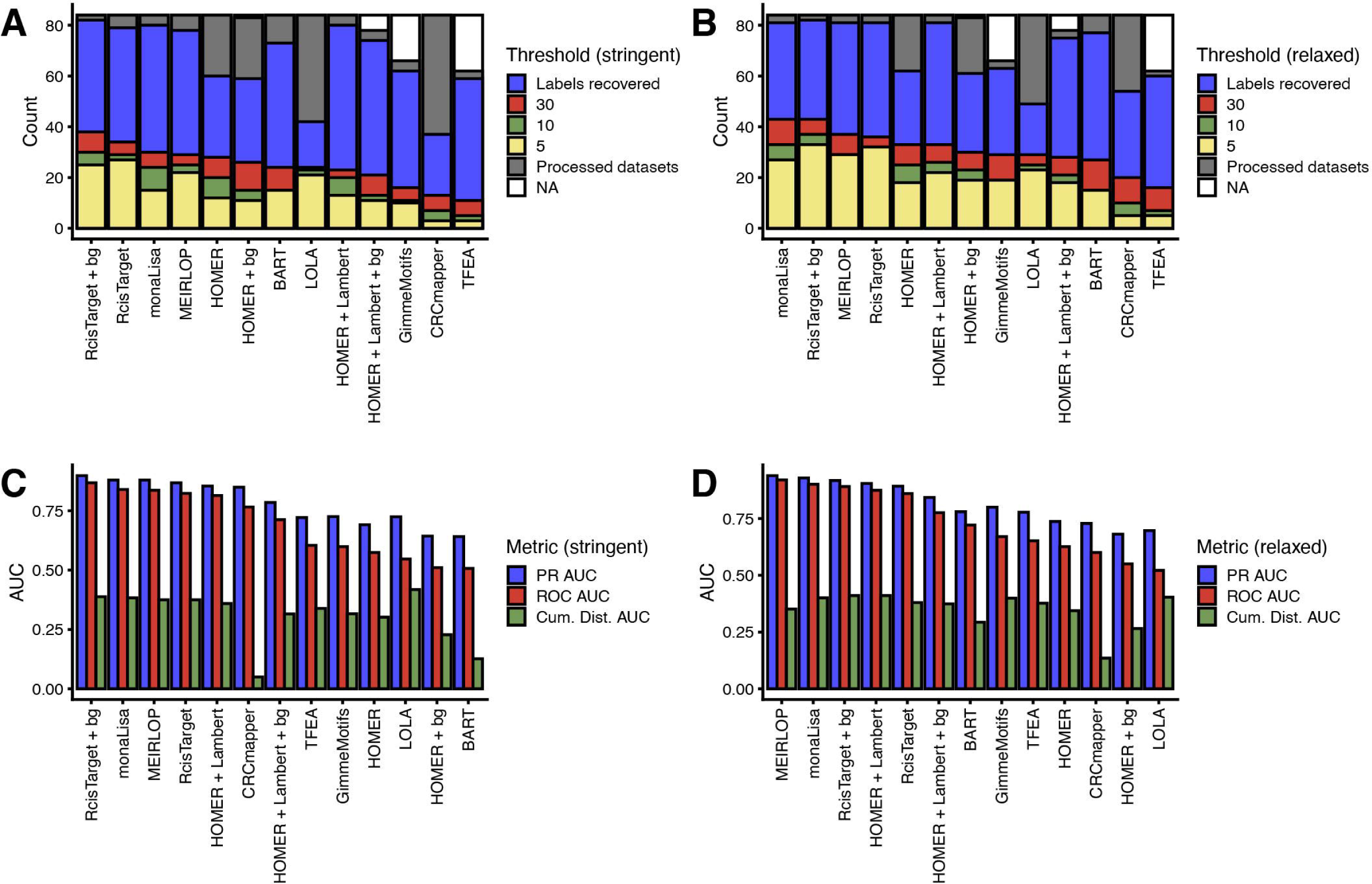
Summary of performance metrics used to evaluate TF prioritization tools. (A) Number of recovered perturbed TFs among the top 5 (yellow), 10 (green), 30 (red) and all ranks (blue) using the stringent label definition. Grey indicates number of successfully processed data sets, but none of the perturbed TF was recovered. White illustrates number of failed data sets. TF prioritization strategies were sorted according to the number of recovered TFs among the top 30. (B) Same as (A) using the relaxed label definition for the recovery of perturbed TFs. (C) Summary of the area under curve (AUC) for precision-recall (PR) curve, receiver operating characteristics (ROC) curve and cumulative rank distribution. (D) Same as (C) but using the relaxed label definition.

The second metric we considered was the number of labels recovered as one of the top 5, 10, or 30 TFs reported in the results (Fig. 2A, B). The rationale behind this metric was based on a plausible real-world scenario that top TFs would often be selected for follow-up experiments. This revealed that RcisTarget, RcisTarget + bg, monaLisa and MEIRLOP were performing best independently of the rank thresholds and label recovery criteria (Fig. 2A, B). In contrast, the bottom ranking tools included GimmeMotifs (stringent), BART (relaxed), CRCmapper and TFEA (both). Nevertheless, even the best performing tools predicted TF labels among the top 30 ranks for only about half of all data sets (e.g., RcisTarget + bg n=43 for relaxed, n=38 for stringent and monaLisa n=43 for relaxed and n=30 for stringent).

We evaluated the tools using the area under the curve (AUC) for the precision-recall curve (PR), receiver operator characteristic curve (ROC), and the cumulative distribution of label ranks. Using these metrics, the best performing tools were again RcisTarget +/- bg, monaLisa and MEIRLOP, independently of the label recovery criteria (Fig. 2C, D and Fig. S4B, C). In the stringent case, the highest AUC of the PR or ROC curves was achieved by RcisTarget + bg (0.90/0.87) and for relaxed by MEIRLOP (0.94/0.92). In contrast, the lowest PR/ROC AUC had BART (0.64/0.51) for stringent label recovery and LOLA (0.70/0.52) for relaxed label recovery. Moreover, the relaxed label recovery criteria led to a slight increase in both metrics for most tools (Fig. S4D, E). CRCmapper and LOLA were the exceptions, showing a decrease in both PR/ROC AUCs.

Finally, the AUC of the cumulative distribution of label ranks confirmed the frontrunner tools mentioned above (Fig. 2C, D and Fig. S4A). CRCmapper was at the bottom of the ranking (0.05/0.14) and BART second last in the stringent evaluation (0.13), while HOMER + bg was second last in the relaxed evaluation (0.27).

In conclusion, the tested tools were able to recover known TF labels with variable accuracies, and monaLisa, RcisTarget and MEIRLOP performed best across several of our benchmark metrics.

### Effects of parameter tweaking on the performance of TF prioritization tools

We next evaluated how modifying the default parameters influenced the performance of the TF prioritization tools. To maintain the number of computational jobs tractable, we selected two or three parameters of each tool based on the emphasis that these parameters were given in the documentation of the tools (see Supplementary Material). We varied the parameters to different degrees, resulting in more than 18,500 computational jobs. For most tools, changing the default parameters had little effect on their overall performance (Fig. S7). For MEIRLOP, however, we observed a drop in performance when varying the default parameters. Importantly, the ranking of TF tools when ran with the default parameters was almost identical to the ranking of TF tools when selecting the runs with the best performing parameters values. Based on these data, we conclude that compared to the choice of the TF tool, varying parameters of an individual tool has a minimal effect in their performance.

### Performance of TF prioritization tools using ATAC-seq data

In addition to H3K27ac maps, 11 of the 84 curated datasets in this study included ATAC-seq maps for 14 TF perturbations and their corresponding controls (Table S1). Using these data, we evaluated how the TF prioritization tool rankings differed when using ATAC-seq instead of H3K27ac maps as inputs. When using ATAC-seq data, the best 3 performing tools to recover the labels among the top 30 hits were monaLisa, MEIRLOP and HOMER, recovering 9, 8, and 8 TF labels, respectively (Fig. S6B).

For most tools, we found that TF labels that were recovered among the top hits using H3K27ac data also ranked among the top hits using the matching ATAC-seq data. For example, out of the 14 TF experiments with both H3K27ac and ATAC-seq, 7 TF labels were recovered by monaLisa among the top 30 hits using H3K27ac data (Fig. S6B, D). Of these, 6 TF labels were also recovered among the top 30 hits using ATAC-seq data. This statistic varied slightly among the different tools (7 out of 7 for BART, 5 out of 5 for HOMER, 5 out of 5 for GimmeMotifs, and 5 out of 6 for LOLA). For BART, the rankings based on ATAC-seq data were identical to the rankings of H3K27ac data. This similarity in the rankings is explained by how BART maps input data into their resource of cis-regulatory elements.

Overall, the best performing tools at identifying TF master regulators from H3K27ac data were also the best tools for ATAC-seq data.

### Influence of experimental and data set features on tool performance

Having established the performance of each tool, we next asked what features could best explain the observed tool performance. To address this question, 16 features were chosen based on the experimental design (e.g., H3K27ac ChIP antibody, perturbation type, etc.), the quality of ChIP-seq (e.g., sequencing depth, fraction of reads in peaks (FRIP), etc.), the effect of the TF perturbation on gene expression as measured by RNA-seq (n=53; e.g., expression of the perturbed TF, etc.; see Methods) and the information content (IC) of the PWM. We trained a random forest classifier using these features to predict the combined stringent and relaxed TF label ranks. Resulting models were able to fit these ranks with a median Pearson correlation coefficient (PCC) between 0.78 (LOLA) and 0.38 (CRCmapper, Fig. 3A).

**Fig. 3.**
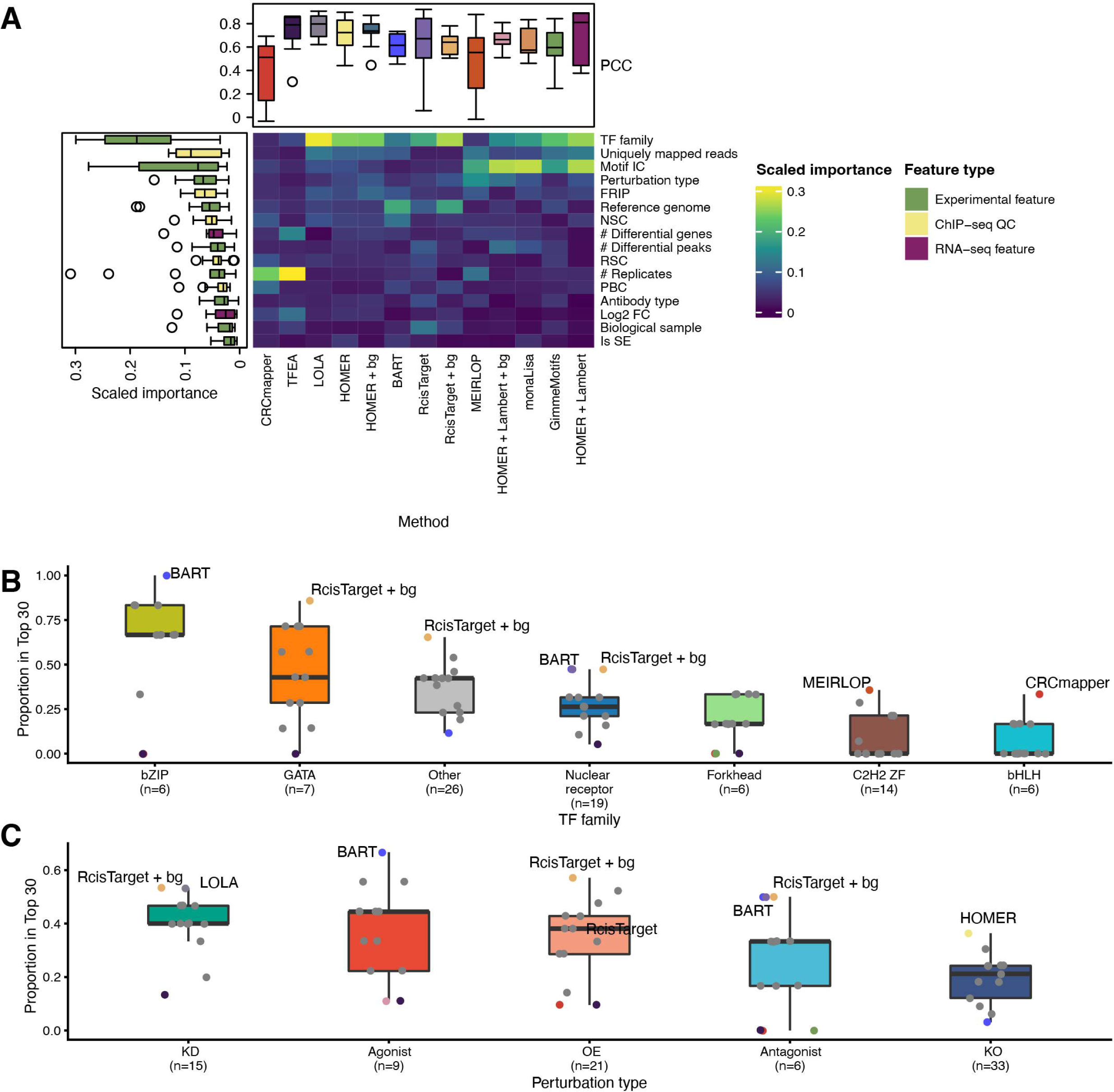
Influence of experiment features on tool performance. (A) Scaled feature importance for 15 features used to regress the TF rankings using random forest models (see Methods). Colour scale encodes the different feature types including experiment (red), ChIP-seq QC (blue) and RNA-seq (green) features. Outlier tools were annotated. Abbreviations: FRIP = Fractions of reads in peaks, Motif IC = Motif information content, NSC = Normalized Strand Cross-correlation coefficient, RSC = Relative Strand Cross-correlation coefficient, PBC = PCR bottleneck coefficient, Is SE = Is Super-Enhancer. (B) Tool performance as proportion recovered TF labels in top 30 stratified by TF families. Selected tools were highlighted. TF families with less than 5 data sets were summarised as “Other”. (C) Same as in (B) but data sets were stratified by perturbation type.

Next, we computed the scaled importance for each feature to gain insights into their influence on resulting rankings (see Methods). This revealed that the most important features were the TF family, the information content of the motif, the number of uniquely mapped reads, FRIP and the type of TF perturbation (Fig. 3A). In contrast, the least important features were the association with a super-enhancer (is SE), RNA-seq log2(FC) of the perturbed TF, biological sample type (tissue vs. cell line) and PCR bottle neck coefficient (PBC).

However, some tools showed deviations from these general patterns. For example, the most important feature for MEIRLOP was the TF perturbation type (Fig. 3A). Other outliers in the feature importance ranking included CRCmapper and TFEA with the number of replicates and BART with the species (reference genome).

Next, we focused on the two most important experimental features and examined whether tools perform differently for each feature modality by looking again at the TF label recovery among the top 30 ranks. Firstly, the overall performance differed across TF families with bZIP and GATA factors being most frequently recovered (e.g., BART bZIP=6/6, RcisTarget bZIP n=5/6 and RcisTarget + bg GATA n=6/7, Fig. 3B). In contrast, TFs belonging to the C2H2 ZF and bHLH families were recovered least frequently (e.g., MEIRLOP C2H2 ZF n=5/14 and CRCmapper bHLH n=2/6). RcisTarget showed the best performance for four out of seven categories across TF families (Fig. 3B). Secondly, the other most important experiment feature was the perturbation type. Overall, we observed a maximum top 30 recovery of 66.7% (Agonist TFs: BART) and a minimum of 0% (Antagonist TFs: CRCmapper, GimmeMotifs, TFEA; Fig. 3C). When considering the median performance, the lowest performance was associated with TF KO experiments (21%, 7/33). RcisTarget performed best in 3 out of the 5 perturbation type categories. Only BART outperformed RcisTarget for agonist perturbations and HOMER for KO (Agonist: 66.7% BART compared with 55.6% RcisTarget; KO: 36.4% HOMER compared with 30.3% RcisTarget + bg).

In summary, TF rankings of the benchmarked tools overall were mostly influenced by TF family and perturbation type, with a tendency of more specialized tools being also influenced by their specific assumptions (e.g., CRCmapper, TFEA).

## Discussion

In this benchmark study, we examined the performance of nine TF prioritization tools in combination with the two most common parameters (PWM motif library and the set of background sequences used), resulting in 13 approaches to rank TFs [16–24]. The ground truth for this was defined using a collection of published H3K27ac ChIP-seq experiments which included a TF perturbation in their design (OE, KO, etc.). The major task for all approaches was to recover these known TF labels.

Figure 4 summarises our results and illustrates the performance of each tool encoded into three groups (poor, intermediate and good) across all considered metrics (see Methods). In our benchmark, we use the default parameters recommended by the tool authors, which were likely selected based on a parameter optimization process during their development. As such, our study also evaluates how generalizable these parameters are across real-world datasets. Thus, a method that performs well across datasets without fine-tuning each parameter is ranked better than a tool that would need dataset-specific fine-tuning of parameters. Nevertheless, our analyses indicate that the effects of tweaking parameters of a tool on its performance is minimal compared to the choice of TF tool.

**Fig. 4.**
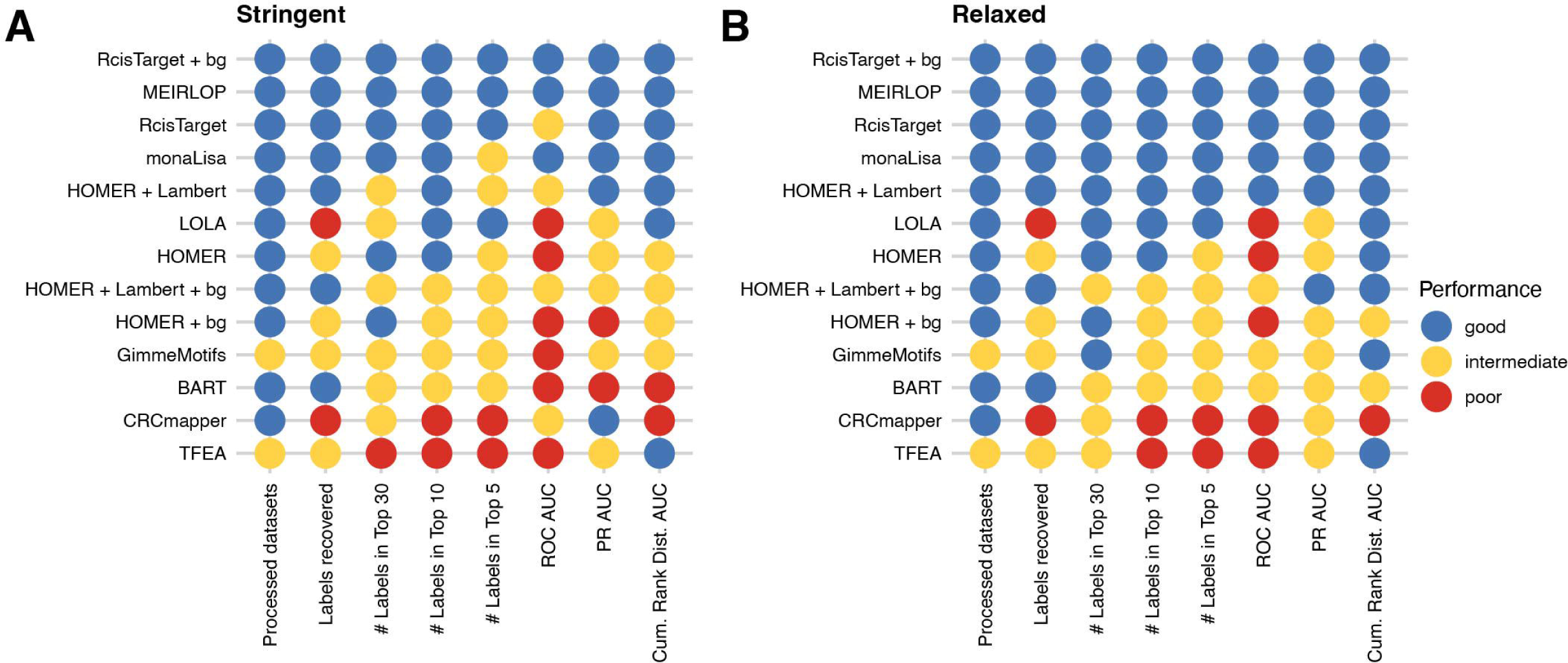
TF prioritization tool benchmark summary. (A) Dot plot heatmap summarising the benchmark results for the stringent label recovery strategy. Tool performance for each single metric was encoded according to the respective rank into one of three categories including poor (blue), intermediate (yellow) and good (red). Tools were ordered according to their overall performance across all eight metrics. (B) Same as (A) but for the relaxed label recovery strategy.

Overall, most tools perform best for the metrics ‘number of successfully processed data sets’ and ‘labels recovered’, suggesting that all tools can process the input data. However, tools showed marked performance differences when considering the other metrics. Based on our performance metrics, we found that RcisTarget and monaLisa perform best, regardless of the TF label recovery criteria used. GimmeMotifs and TFEA failed to complete for around 20% of the test datasets, but their label recovery was relatively good when these tools ran thought successfully. Our analysis thus indicates that these tools could substantially boost their performance by increasing the robustness of their code implementation.

We found that all TF prioritization approaches perform better using the relaxed label recovery criterion (see Results). Moreover, differences between the stringent and relaxed label recovery criteria were only observable for tools in the bottom half of the final rankings. Top ranking approaches like RcisTarget, MEIRLOP and monaLisa already performed well using the more stringent criteria. In contrast, approaches in the bottom half profited from the relaxed criterion due to the circumstance that they ranked a homologous TF even better than the exact TF label.

Furthermore, we investigated the influence of pre-defined genomic background sequences (‘+bg’) and/or the use of a more comprehensive consensus motif library (Lambert *et al.*) if tools were enabling the user to specify these parameters [2]. This revealed that for example RcisTarget profited from specifying a custom background, but HOMER worked better using its default background computation. In contrast, HOMER performed better using the Lambert *et al.* motif library instead of the default one. Although the choice of background seemed to partially influence the performance of the tools, this was neglectable compared to the choice of TF ranking tool or other parameters such as the motif library.

The bottom three tools were TFEA, CRCmapper and BART, performing either ‘poor’ or ‘intermediate’ across most metrics. The poor performance of TF ChIP-seq library-based approaches such as BART might be attributed to a lower complexity of their underlying databases compared with PWM-based tools. Since the enrichment approaches of BART and RcisTarget are quite similar, one could speculate that the incorporation of large-scale TF datasets such as REMAP 2022 or UNIBIND could greatly enhance the performance of such tools [25,26]. In contrast, the poor performance of CRCmapper could be explained by the specific assumptions made by the tool: CRC is optimized for recovering TFs in SEs, and thus expects that TFs of interest are associated with a SE, which might not broadly apply across multiple experiments and datasets [20]. Overall, we observed for 25 out of 84 H3K27ac data sets an association of the perturbed TF to a SE. Therefore, CRCmapper’s very specific assumptions led to an overall poorer performance in our benchmark, which focused on a more general task.

We found that the families of the TF substantially influence the recovery of the TFs from the tools. This observation is in line with previous reports of varying performance of PWMs to predict TF binding depending on their TF family affiliation (e.g., C2H2 ZFs and bHLH TFs) [27]. The tools benchmarked in this manuscript depend on PWMs, and thus their performance could be compromised when PWMs are not sufficient to accurately predict TF binding to DNA. As an alternative to PWM-based methods, deep learning approaches have recently been developed to predict TF binding. For example, DeepBind and BindSpace are convolutional neural network models developed to predict transcription factor binding [28,29]. Another recent development is the Enformer model, that was able to predict dozens of chromatin and gene expression tracks uniquely from DNA sequence [30]. A major advantage of these models is their capacity to learn not only motifs, but also sequence features such as DNA sequence composition and complex positional configurations, such as periodicity of TF motifs or distance requirement between TF motifs. Our benchmark suggests that tools to prioritize TFs would benefit from incorporating deep leaning-centric predictions of TF binding (for an in-depth discussion of TF binding prediction models see [31]).

Although we compiled a large H3K27ac dataset for our benchmark, this study has some limitations. First, we benchmarked these tools on H3K27ac ChIP-seq data, assuming that the TF perturbation will lead to H3K27ac changes. Future work is needed to evaluate their performance using other high throughput sequencing technologies, such as H3K4me3 (Promoter), H3K4me1 (Enhancer) ChIP-seq, RNA-seq and a more comprehensive ATAC-seq (open chromatin) dataset collection [1]. Second, this benchmark is focused on the performance of approaches to recover a perturbed TF, mimicking a particular real-world scenario common, for example, in drug discovery. As such, we do not assess the performance of the tools in other contexts (e.g., simulation approaches, other definitions of regulatory elements such as open chromatin, or other histone marks).

## Conclusion

In conclusion, our comprehensive benchmark provides recommendations for the scientific community on which TF prioritization tool perform best (i.e., RcisTarget, MEIRLOP and monaLisa) for perturbed TF recovery. We believe this will help improve hypothesis generation from H3K27ac ChIP-seq data, one of the most widely profiled histone marks. In addition, our study reveals shortcomings of current tools, which we are hoping will influence further improvement of existing tools as well as the development of novel tools.

**Fig. S1.**
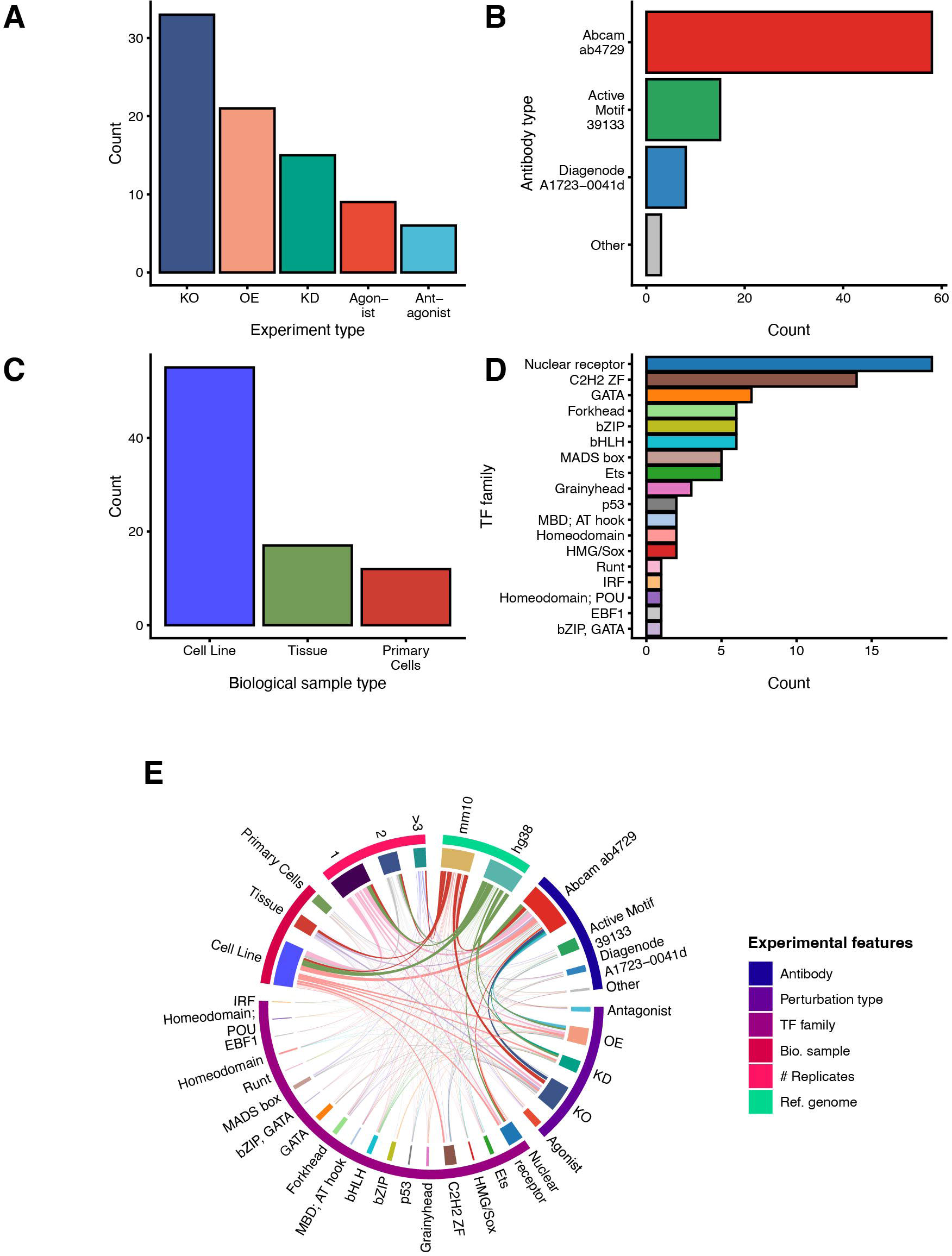
Overview of the manually curated H3K27ac data sets with underlying TF perturbation. (A) Barplot illustrating the number of ChIP-seq data sets across different TF perturbation categories. (B) Number of data sets stratified by the H3K27ac antibody used for the ChIP. (C) Number of H3K27ac data sets split by the biological sample type. (D) Number of H3K27ac data sets stratified by the TF family of the perturbed TF. (E) Circos plot displaying the cross dependencies of the different categorial variables across all 84 H3K27ac ChIP-seq data sets. Links were scaled by the frequency of variable co-occurrence.

**Fig. S2.**
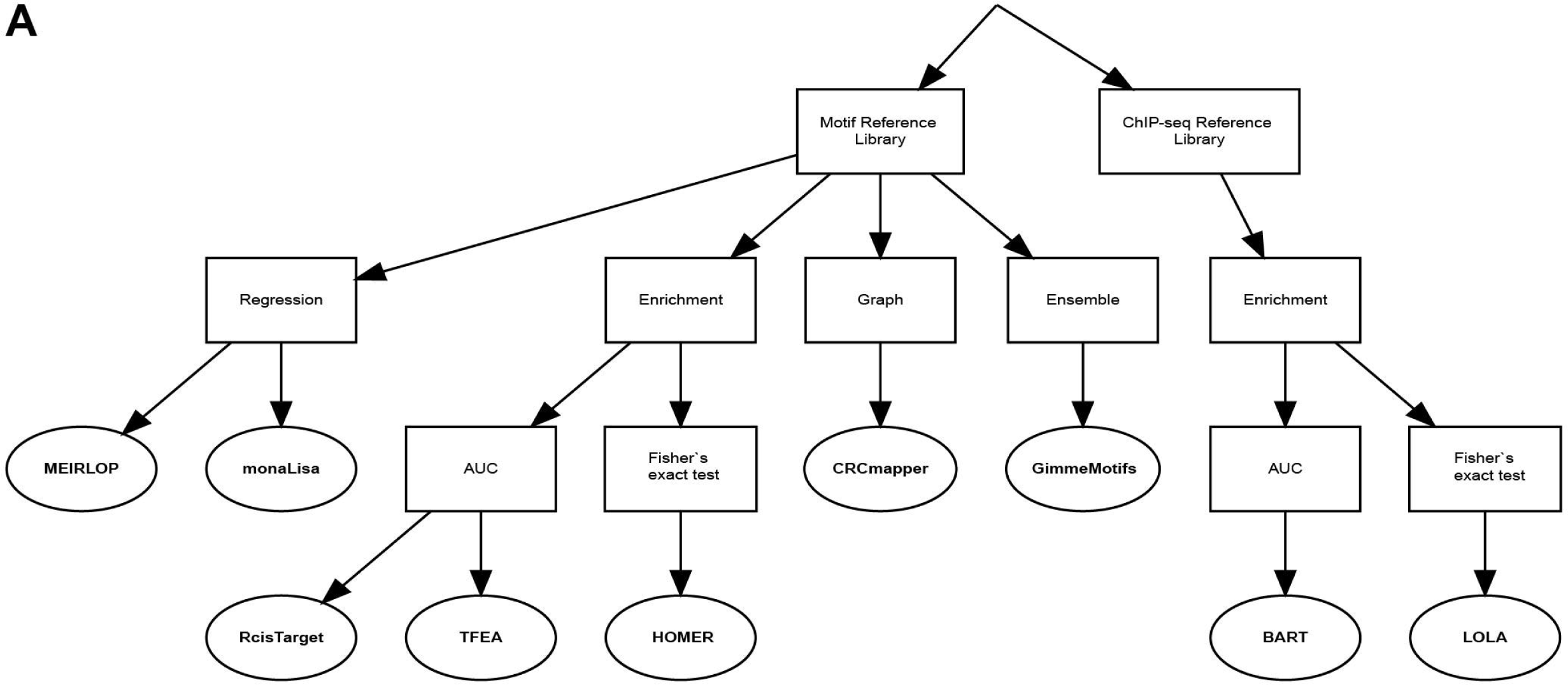
Overview and classification of the selected TF prioritization tools examined in this benchmark study. For detailed description of tools see Supplementary Methods.

**Fig. S3.**
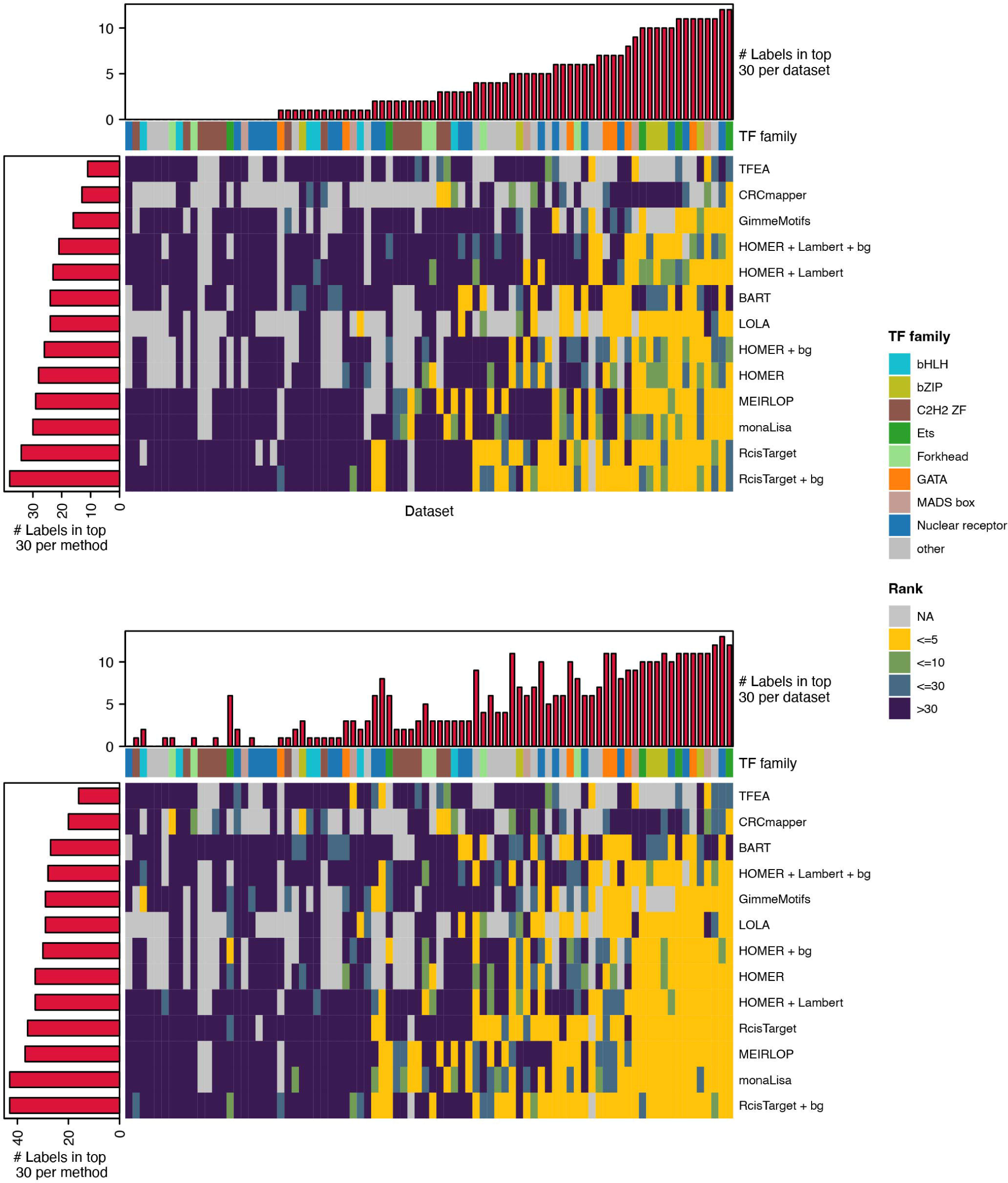
Overview of TF label recovery for each data set and tool. (A) Heatmap illustrating the TF label recovery using the stringent label definition, across data sets and per tool, among the top 5 (yellow), 10 (green), 30 (cyan), in the entire ranking (dark blue) or not being included/failed run (grey). Row barplot shows the number of recovered TF labels among the top 30 for each tool. Column barplot shows the number of tools recovering a particular TF label in their top 30 ranks. (B) Same as (A) but for the relaxed TF label definition.

**Fig. S4.**
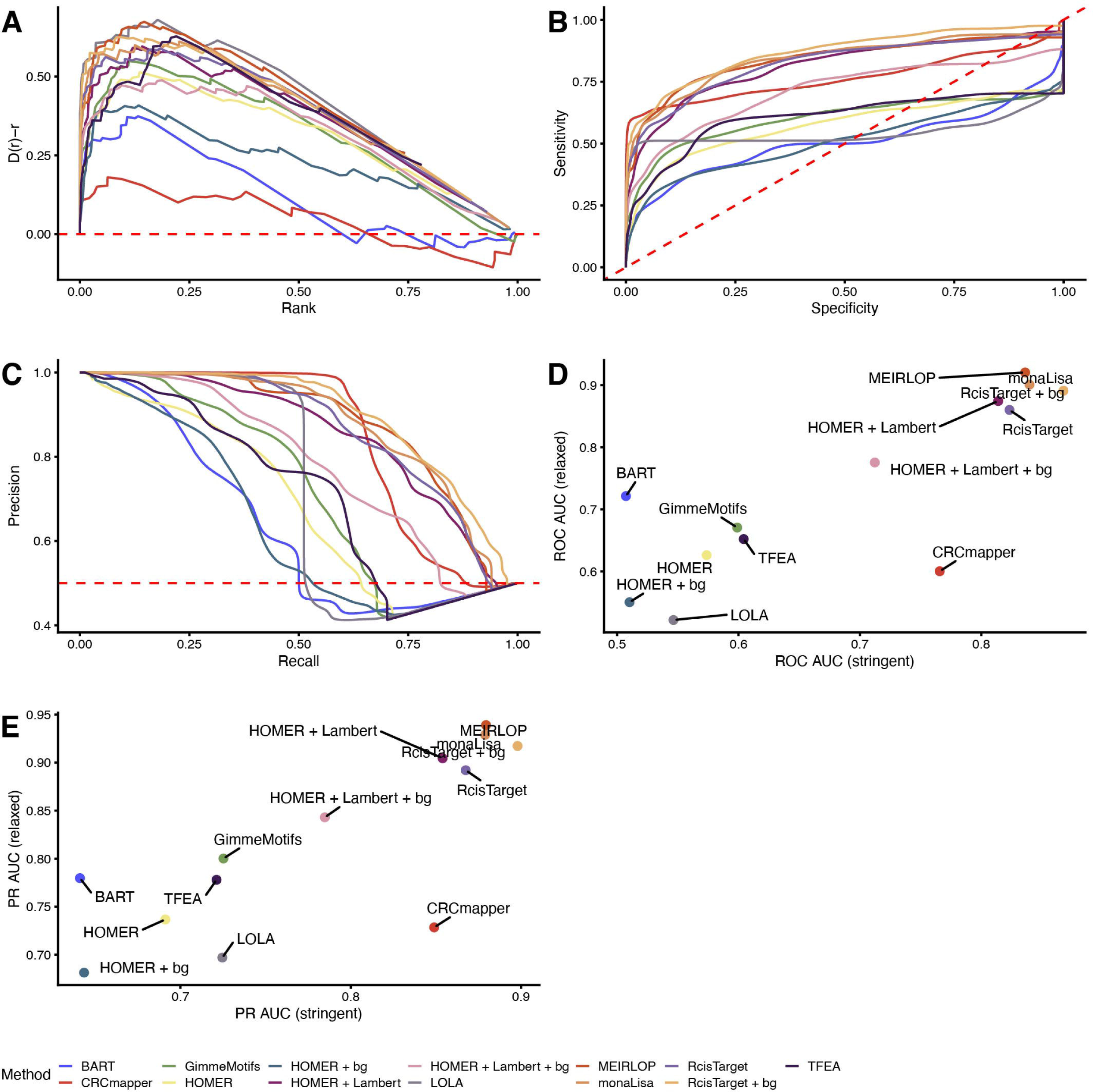
Overview and comparison of AUC based performance metrics. (A) Cumulative distribution of scaled ranks for each TF prioritization tool (stringent label recovery). (B) Average ROC curves, per TF prioritization tool, over 5,000 bootstraps using the stringent label recovery. (C) Average PR curve, per TF prioritization tool, over 5,000 bootstraps using the stringent label recovery. (D) Scatterplot comparison of ROC AUCs between stringent and relaxed label definition. (E) Scatterplot comparison of PR AUCs between stringent and relaxed label definition.

**Fig. S5.**
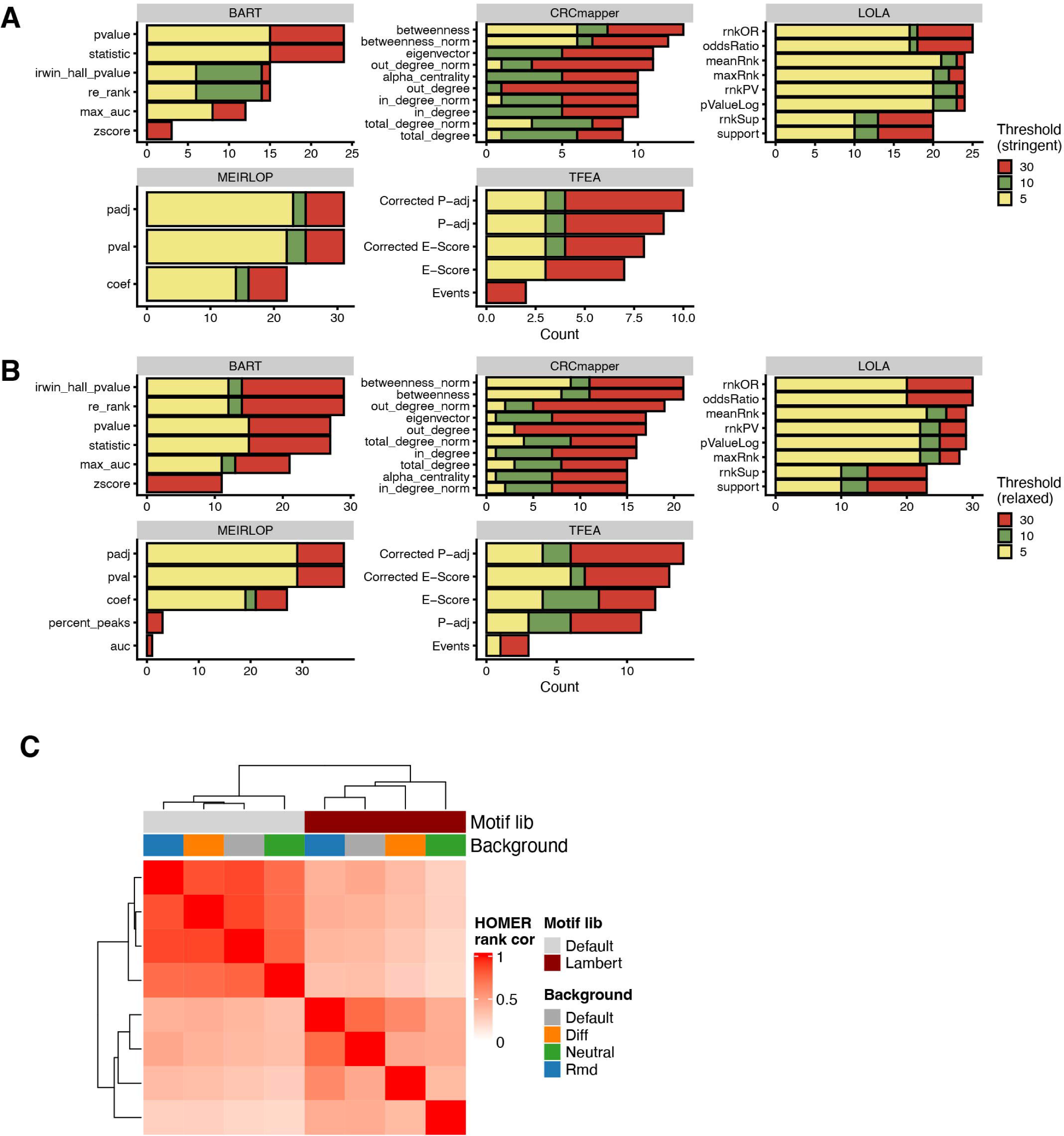
Comparison of ranking metrics and backgrounds for TF prioritization. (A) Number of recovered perturbed TFs among the top 5 (yellow), 10 (green) and 30 (red) using the stringent label definition for each ranking metric outputted by a tool (panels). (B) Same as (A) using the relaxed label definition for the recovery of perturbed TFs. (C) Heatmap of the Pearson correlation coefficients between the rankings of TF labels across the 84 ChIP-seq data from 8 different HOMER setups. Top annotation illustrates used parameters: 1) Motif library: HOMER default (grey) or Lamber *et al* (red) and 2) different background sets: default (grey), differential (orange), neutral (green) and random (blue; see Methods “Background comparison”).

**Fig. S6.**
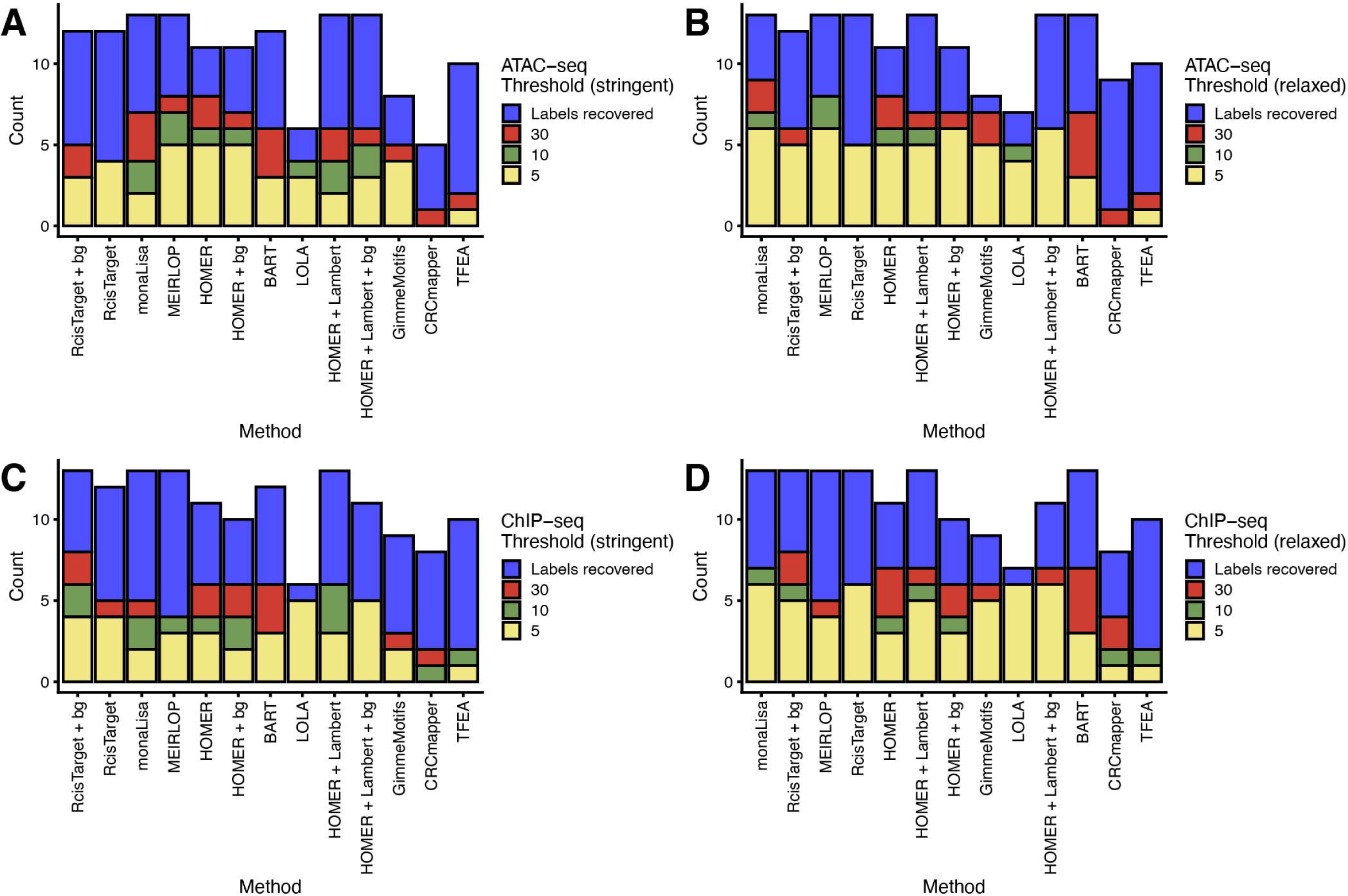
TF prioritization tool evaluation based on matched ATAC-seq samples. (A) Number of recovered perturbed TFs among the top 5 (yellow), 10 (green), 30 (red) and all ranks (blue) using the stringent label definition. TF prioritization strategies were sorted according to Barplot in Fig. 2. (B) Same as (A) using the relaxed label definition for the recovery of perturbed TFs.

**Fig. S7.**
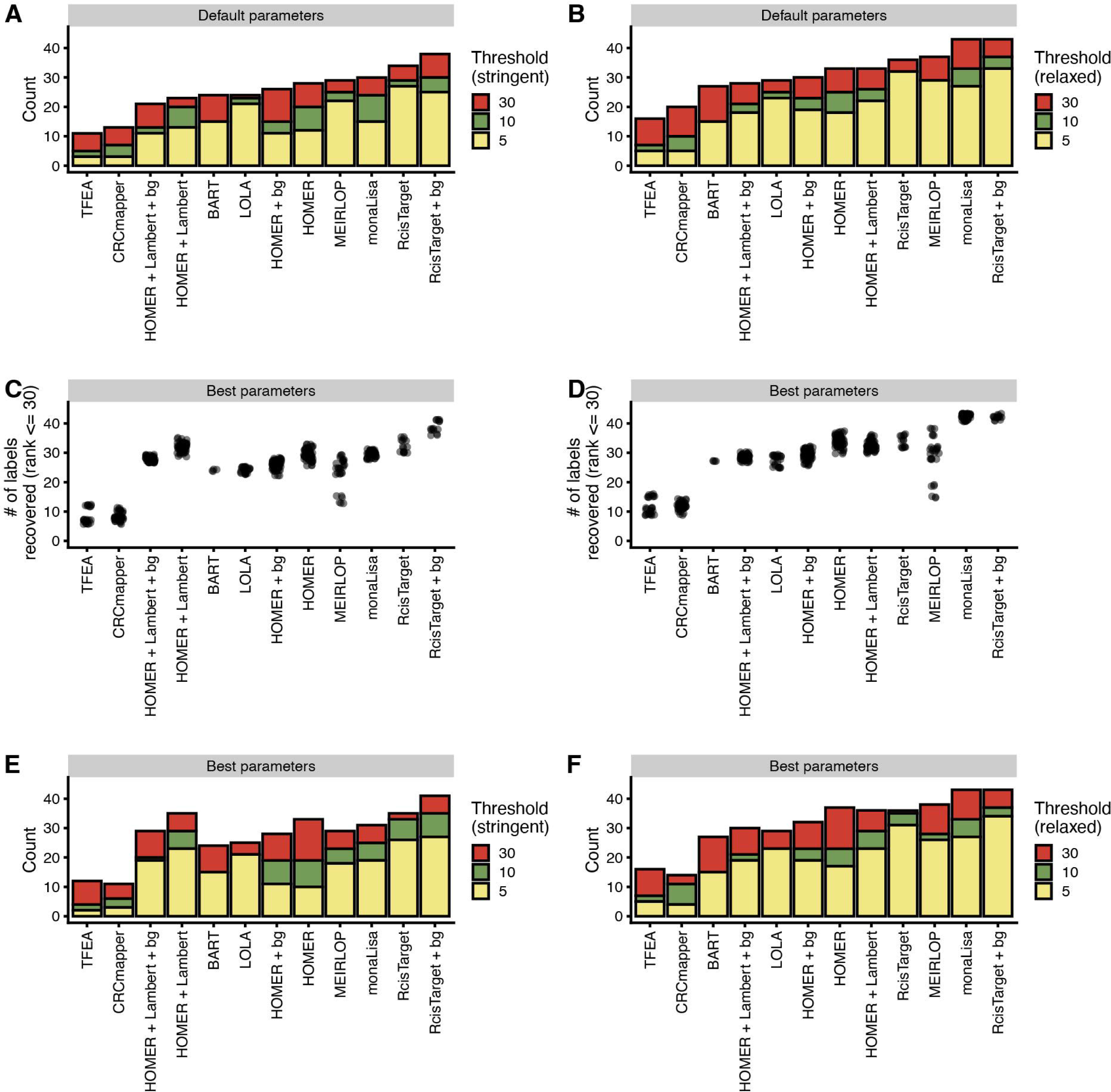
Comparison of different parameters for TF prioritization. (A) Number of recovered perturbed TFs among the top 5 (yellow), 10 (green) and 30 (red) using the default parameter setting and stringent label definition. (B) same as (A) using the relaxed label definition for the recovery of perturbed TFs. (C) Number of recovered perturbed TFs among the top 30 for different parameter settings and stringent label definition. Plot sorted according to (A). (D) same as (C) using the relaxed label definition. Plot sorted according to (B). (E) same as (A) using the parameter set maximising the number of TF labels recovered among the top 30. (F) Same as in (E) using the relaxed label definition.

**Table S1.** Table summarising 84 H3K27ac ChIP-seq experiments with TF perturbations.

**Table S2.** List of the TF prioritization tools benchmarked in this study.

## Methods

### ChIP-seq pre-processing

Publicly available H3K27ac ChIP-seq data sets with TF perturbations were manually curated and associated FASTQ files were downloaded using SRAToolkit (V2.11.2; https://github.com/ncbi/sra-tools). Pre-processing of each experiment was performed using the ENCODE ChIP-seq pipeline (V1.9.0) [32]. Briefly, reads were aligned to the respective reference genome (hg38 or mm10) using Bowtie2 (V2.3.4.3) and subsequently filtered for unmapped reads, not primary alignments as well as duplicates using SAMtools (V1.12)/Picard (V2.9.2) [33–35]. Peak calling was performed using MACS2 (V2.2.5) with following parameters: --cap-num-peak 500000 --pval-thresh 0.01 [36]. Consensus peak sets per condition were computed by performing the overlap reproducibility analysis as implemented in the ENCODE pipeline. In addition, peaks were filtered for overlap with blacklist regions.

### ATAC-seq pre-processing

We scanned the publicly available ChIP-seq data (see above) for matched ATAC-seq data sets with TF perturbations. The associated FASTQ files were downloaded using SRAToolkit (V2.11.2; https://github.com/ncbi/sra-tools). Pre-processing of each experiment was performed using the ENCODE ATAC-seq pipeline (V2.0.3) [32]. Briefly, reads were aligned to the respective reference genome (hg38 or mm10) using Bowtie2 (V2.3.4.3) and subsequently filtered for unmapped reads, not primary alignments as well as duplicates using SAMtools (V1.12)/Picard (V2.9.2) [33–35]. Peak calling was performed using MACS2 (V2.2.5) with following parameters: --cap-num-peak 300000 --pval-thresh 0.01 [36].

Consensus peak sets per condition were computed by performing the overlap reproducibility analysis as implemented in the ENCODE pipeline. In addition, peaks were filtered for overlap with blacklist regions.

### Differential peak analysis

For the differential peak calling, peaks from both conditions (control and TF perturbation) were merged. H3K27ac as well as open chromatin (ATAC-seq) enrichment was quantified for these merged peaks, by counting the reads using the featureCount function from the Rsubread package (V2.2.6) with parameters countMultiMappingReads = False and allowMultiOverlap = True [37].

For experiments with more than one replicate per condition, differential peak analysis was performed using DEseq2 (V1.30.1) with default settings [38]. All peaks were then sorted by - log10(p-value) * log2(fold change) (log2(FC)) and we took the top 1,000 peaks as foreground and the bottom 1,000 peaks as background set.

For experiments with only one replicate per condition, we normalized the counts using DESeq2 estimateSizeFactors function and calculated the log2(FC). Peaks were sorted according to their log2(FC). The top 1,000 peaks were defined as foreground and the bottom 1,000 as background sets.

Resulting foreground were used as input for TF prioritization tools expecting peaks as input (e.g., HOMER, RcisTarget, etc.). In case of the “+ bg” strategy, we provided the background peak sets as custom background.

### RNA-seq pre-processing

RNA-seq data associated with the H3K27ac ChIP-seq was downloaded using SRAToolkit. Expression levels for the respective gene annotation (Ensembl GRCm38.98 or GRCh38.98) was performed using the PISCES pipeline (V0.1.3.1) with default parameters [39,40].

### Differential gene expression analysis

The function getBM from the package biomaRt (V2.46.3) was used to assign the external_gene_name to the ensembl_gene_id from Ensembl [40]. We then used DEseq2 to normalize the raw gene counts and fit them to a negative binomial distribution. Then a generalized linear model and Wald test was used to compute differential expression between the TF perturbation condition compared with the control [38].

### TF prioritization tool settings and parametrizations

A comprehensive tool overview including versions can be found in Supplementary Table 2.

#### 1) BART

For this benchmark, BART was run with the positional parameter ‘region’ using the differentially expressed genomic region sets described in differential peak analysis as input [17]. The output of BART was ranked according to the p-value column.

#### 2) CRCmapper

For this benchmark, we computed potential SEs using ROSE2 separately for condition and control. We then ran CRCmapper on both sets of .bam files, identified peaks (see ChIP-seq pre-processing), activity tables, and the default parameters of CRCmapper [20]. To infer differentially expressed TF’s, we computed the normalized output degrees individually from condition and control CRCmapper outputs as a summary network statistic. Finally, the differential network statistics were calculated as the difference between condition and control betweenness and were subsequently used for ranking.

#### 3) GimmeMotifs

GimmeMotifs was run using its gimme maelstrom command and its second input option which contains the merged peaks from control and condition experiment identified in ChIP-seq pre-processing step and their log-transformed read counts [24]. The reference library used is the Lambert *et al.* motif library. We also allowed the tool to return redundant motifs, to report the scores of all motifs and use 12 threads by using the parameters: --no-filter, – filter_cutoff 0 and –N 12. All other parameters were left at their default values. The output of GimmeMotifs used for ranking was z-scores.

#### 4) HOMER

We ran HOMER four times for our benchmark: Once using HOMER’s default motif library and using no background sequences but instead letting HOMER select them from the input, once using HOMER’s default motif library and using background sequences as computed in differential peak analysis (HOMER + bg), once using the Lambert *et al.* motif library as a reference library and no background sequences (HOMER + Lambert) and finally using the Lambert *et al.* motif library as a reference library and using the pre-computed background sequences (HOMER + Lambert + bg) [2,16]. As input sequences we always used the differentially expressed peaks as computed in differential peak analysis. HOMER’s script findMotifsGenome.pl was ran with the above descript parameters and inputs, as well as the parameter –nomotif to indicate that we are not interested in de novo motif enrichment. All other parameters were left to their default values. HOMER’s output used for ranking were the p-values.

#### 5) LOLA

For our benchmark, we ran LOLA with the query set being the differentially expressed peaks as discussed in differential peak analysis [18]. The universe or background peaks used are the combined peaks from the condition and control experiment computed as in ChIP-seq pre-processing. LOLA was then run with its default parameters and using its default reference library of public datasets. For ranking we used LOLA’s mean rank based on p-value, log odds ratio and number of overlapping regions.

#### 6) MEIRLOP

In our benchmark, we used fasta files containing the merged peaks from control and condition experiment identified in ChIP-seq pre-processing step and their associated log2(FC) (see differential peak analysis) as scores for the input of MEIRLOP [21]. The Lambert *et al.* motif library was used as the reference library and the –length parameter was set to incorporate sequence length as a covariate since our input sequences were not of the same length as is preferred by MEIRLOP. All other parameters were left to their default values. We ranked the output of MEIRLOP according to the output’s adjusted p-value.

#### 7) monaLisa

To run monaLisa we used its randomized lasso stability selection on our precomputed differentially expressed regions (see differential peak analysis) with the response vector corresponding to their log2(FC) [22]. As predictors the Lambert *et al.* motif library was used. All other parameters were kept at the same values as indicated in their vignette. MonaLisa’s output was ranked according to the normalized area under the selection curve.

#### 8) RcisTarget

We ran RcisTarget twice: Once using the differentially expressed peak regions (see differential peak analysis) with (RcisTarget + bg) and once without background regions (RcisTarget) [19]. The background regions are the merged peaks from control and condition experiment identified in the ChIP-seq pre-processing step. We set the NES threshold parameter to 0, such that all motifs are returned even if the predicted NES score is very low. All other parameters were set as suggested by the vignette on ‘RcisTarget - on regions’. The output of RcisTarget was ranked according to the NES score.

#### 9) TFEA

To run TFEA, we used the BAM and BED files of the control and condition experiments as computed in ChIP-seq pre-processing and the Lambert *et al.* [2,23]. Motif library. TFEA was then ran in parallel with the parameter –cpus 6 and all other parameters set to the default values. TFEA’s output was ranked according to the Bonferroni and GC corrected p-values.

### Performance benchmark

Depending on the approach, the outputs contain either a list of TF or motifs, with associated scores attached. To account for different types of scores reported by the approaches (e.g., p-value, z-score, AUC, …), we ranked the entries in the outputs according to their score, with lower ranks associated with more important entries. We then compared the ranks of labels scaled between [0, 1].

Two label identification strategies were employed to account for the advantage of approaches using reference libraries which allow motifs to be associated with multiple TFs (Figure 2A). The first strategy termed ‘stringent’, forces a one-to-one mapping between motifs and TFs in all approaches. The second strategy called ‘relaxed’ does the exact opposite by creating a mapping between each TF and TFs it is sharing a motif with within the Lambert *et al*. motif library. The rank of the label is then computed by using the best rank between the mapped TF of the label.

We adapted part of our benchmark metrics from the ChEA3 paper published by Keenan *et al.* in 2019 [41]. Accordingly, we calculated a Receiver Operator Characteristic (ROC) and Precision Recall (PR) curve by bootstrapping the down sampled negative class from the rankings as was suggested by Keenan *et al*. By doing this we account for the fact that our positive class consisting of our labels is significantly smaller than our negative class comprised of all other TFs. For both, the ROC and PR curve, we also computed the Area Under the Curves (AUC).

The second metric we implemented from the Keenan *et al.* looks at the deviation of the cumulative distribution of perturbed TF ranks D(r) from a uniform distribution using the Anderson-Darling test. We would expect a significant p-value if perturbed TFs would display preferentially low or high ranks. Additionally, we determined the AUC of D(r)-r since many labels with low ranks give rise to a high AUC in this case.

To put all results together we created a summary of all metric outcomes. We stratified all outcomes into three groups for better readability. For the number of labels recovered and successfully processed datasets the thresholds are determined by dividing all used datasets (84) into three equal groups. The same idea was used for the AUC of ROC and PR curve, where we grouped the results into three groups between 0.5 and 1 and for the cumulative rank distribution AUC between 0 and 0.5. For the number of labels in Top 5-30, we used the maximal respective value to create the three groups.

### Ranking metric comparison

We compared the choice of ranking metric for tools outputting more than two non-correlated metrics which could be used for ranking the TF motifs. Following tools fulfilled these criteria: BART, CRCmapper, LOLA, MEIRLOP and TFEA (Table S2). The H3K27ac analysis using these tools was performed as described above. Resulting outputs were used to create a TF ranking for each metric and count the number of recovered TF labels in the top 30 (Fig. S5A, B). The metric maximizing the number of recovered TFs among the top 30 was used for the benchmark comparison of tools (see Table S2 for list of the best metric per tool).

### Background comparison

To compare the influence of background choice on the TF ranking, we ran HOMER with four different background sets with either the default HOMER motif library or Lambert *et al*. The four different background sets were constructed as following:

- Default – random selection of GC% content matched regions from the genome.
- Diff – Top 1,000 most differential peaks for the control condition (see section differential peak analysis)
- Neutral – 1,000 non-differential peaks from the comparison TF perturbation vs control.
- Rmd – random draw of 1,000 GC% content matched non-overlapping regions from the genome.

HOMER and downstream analysis were performed as described above.

### Parameter tweaking

We conducted a parameter tweaking analysis to ensure the robustness of the final tool ranking. We selected up to three parameters per tool from a set of available parameters based on their potential impact on the results (see Supplementary Text for the full list). We explored up to five different values per parameter in combination with each other.

We performed TF prioritization for each tool with the added parameters, as described in each tool section. The downstream benchmark analysis was conducted as described above.

### Random forest modelling and feature importance

To study the influence of experimental features and data characteristics on TF rankings, we looked at three different types of features: ChIP-seq quality measures (Normalized Strand Cross-correlation coefficient (NSC), uniquely mapped reads, Relative Strand Cross-correlation coefficient (RSC), PBC, Fraction of reads in peaks(FRIP)), experimental features (perturbation type, number of replicates, TF family, antibody type, reference genome, is super-enhancer, biological sample type, number of differential peaks) and RNA-seq features (TF log2(FC), number of differential genes). All groups with less than five experiments were grouped together into ‘Others’.

For RNA-seq features, TF log2(FC) were categorized into four groups (log2(FC) <= 2, 2 < log2(FC) <= 6, log2(FC) > 6 and missing RNA-seq) and the number of differentially expressed genes into five groups (# differential genes <= 100, 100 < number of differential genes <= 500, 500 < number of differential genes <= 1000, number of differential genes > 1000 and missing RNA-seq).

In addition, we computed the information content (IC) for each TF as additional feature. Briefly, the IC per motif was computed using ggseqlogo to compute the IC per bp and then average over each position [42]. From that we derived an IC per TF by averaging the IC per motif over multiple motifs as assigned by Lambert *et al*.

We added all missing experiments with rank=1 to have 84 experiment results for each tool. We trained a random forest with 10-fold cross validation predicting the rank for the above-described features using the cforest() function from the R package *party* (V1.3) [43]. The conditional feature importance was calculated using the varimp() function for each fold. We scaled resulting feature importance and computed the mean across the 10 folds.

Additionally, we calculated the Pearson’s correlation coefficient between predicted and true rank in the test sets to assess the fit of the random forest for each tool.

## Supporting information

Table S1 Table summarising 84 H3K27ac ChIP-seq experiments with TF perturbations.

Table S2 List of the TF prioritization tools benchmarked in this study.

## Ethics approval and consent to participate

Not applicable

## Consent for publication

Not applicable

## Availability of data and materials

All ChIP-seq data sets used in this study were listed in Supplementary Table S1 including GEO IDs.

All intermediate results (peak files, tool outputs and processed outputs) were made available here https://zenodo.org/records/10990183.

Code to reproduce our findings can be found here https://github.com/Novartis/TF_Prioritization_Benchmark_GB2023.

## Competing interests

All authors are, or were, employees or affiliates of the Novartis Pharma AG. The authors declare that they have no competing interests.

## Funding

Not applicable

## Authors’ contributions

SS conceptualized the initial study; SG, CK, SH and EF acquired funding; SS and SG curated the data; SS and LSS designed the methodology; All authors pursued the study conceptualization and contributed to the methodology; LSS conducted the formal analysis; SS supervised the analysis; SS and LSS wrote the original draft; AR, SH, EF, CK, SG and SS revised and edited the manuscript (using https://credit.niso.org).

## Acknowledgments

We would like to thank Dr. Julianne Perner from the Novartis Institutes for Biomedical Research (Basel, Switzerland) for her scientific and technical contributions to this project. We would also like to express our gratitude to Drs. Mikhail Pachkov and Erik van Nimwegen from the Biozentrum of the University of Basel (Switzerland) for fruitful discussions.

## Notes

https://zenodo.org/records/10990183

