## Supplementary text for "Benchmarking tools for transcription factor prioritization"

1) BART

BART uses a curated comprehensive list of millions of MNase hypersensitive sites in the genome, each of which is considered a cis-regulatory element, and a reference catalogue with hundreds of TF binding profiles. For each TF binding profile in the reference catalogue, it annotates each cis-regulatory element according to whether the TF is bound or not. For a query dataset (e.g., a ChIP-seq sample), BART tabulates the number of reads mapping to each cis-regulatory element and ranks the cis-regulatory elements according to the number of read counts. For each transcription factor, it calculates a ROC curve using the cis-regulatory elements ranked according to the signal on the query sample and the TF binding information from the catalogue as gold truth. It then estimates the area under the ROC curve (AUC) and performs a Wilcoxon-test between the AUC values of a given TF versus the AUC values of all other TF factors. It also estimates a background model by calculating the AUC values for H3K27ac profiles and estimates z-scores of each TF with respect to this background AUC distribution. Predictions of transcription factors are then based on the p-value of the Wilcoxon test, the z-scores, and the AUC values.

Reference: *Wang et al.* BART: a transcription factor prediction tool with query gene sets or epigenomic profiles. **Bioinformatics**, 34(16), 2018, 2867–2869.

Software link: <https://zanglab.github.io/bart/index.htm>

2) CRCmapper

The central hypothesis of CRCmapper states that core TFs are driven by super-enhancers (SE) and collectively regulate their own gene expression, resulting in an interconnected auto-regulatory loop. Thus, CRCmapper aims to identify networks of TF that both auto-regulate their own expression and that regulate the expression of other TF. Using chromatin profiles as input signal (e.g., H3K27ac ChIP-seq), CRCmapper identifies super-enhancers located nearby genes coding for transcription factors. CRCmapper does a motif search for all TFs within the super-enhancer elements. Then, it identifies TFs that regulate their own expression by binding to a near super-enhancer, based on the presence and number of TF binding motifs. Using the presence and number of motifs, it then identifies networks of auto-regulated TFs that regulate their expression among each other.

Reference: *Saint-André et al.* Models of human core transcriptional regulatory circuitries. **Genome Res.** 2016 Mar;26(3):385-96.

Software link: <https://github.com/younglab/CRCmapper>

3) GimmeMotifs

The maelstrom algorithm from GimmeMotifs uses a test statistic aggregation approach to rank transcription factors. First, it inputs a matrix of signal per peak and calculates different test statistics for differential motif enrichment between conditions (in our benchmark, perturbation vs control). These include, among others, hypergeometric tests, non-parametric Mann-Whitney U tests and several regression approaches for variable selection. In a second step, it aggregates the ranks of the different test statistics and converts them into z-scores. The z-scores from the different methods are then aggregated using Stouffer’s method into a single z-score. This aggregated z-score is used to prioritise TFs.

Reference: *Bruse et al.* GimmeMotifs: an analysis framework for transcription factor motif analysis. **BioRxiv,** 2018. Doi.org/10.1101/474403

Software link: <https://gimmemotifs.readthedocs.io/en/master/index.html>

4) HOMER

HOMER counts, for each TF, the number of motifs in input targets (e.g., peaks) and background regions. For each TF, it performs a hypergeometric test that test whether there is an over-representation of motifs in the target sequences compared to the background sequences. To avoid spurious enrichments driven by biases in sequence composition, HOMER selects background sequences such that they match the GC content distribution of the input target sequences.

Reference: *Duttke et al.* Identification and dynamic quantification of regulatory elements using total RNA. **Genome Res.** 2019 Nov; 29(11): 1836–1846.

Software link: <http://homer.ucsd.edu/homer/motif/>

5) LOLA

LOLA borrows the methodology from classical gene-set enrichment analyses. LOLA compares the query regions of the genome to a comprehensive catalogue of TF binding sites collected from chromatin profiling experiments and other data sources. For each TF site in the reference catalogue, LOLA calculates a contingency table based on the overlap of the reference TF binding regions with the query regions and with background regions. It performs a fisher exact test to determine whether there is a significant overlap between the query regions and the TF binding regions. LOLA ranks are based on p-value, odds ratio, and number of overlapping regions.

Reference: *Sheffield et al.* LOLA: enrichment analysis for genomic region sets and regulatory elements in R and Bioconductor. **Bioinformatics,** 2016 Feb; 32 (4), 587–589.

Software link: <https://www.ncbi.nlm.nih.gov/pmc/articles/PMC4743627/>

6) MEIRLOP

MEIRLOP uses logistic regressions to determine whether the intensity of an input signal is associated to the presence of a motif within a sequence. First, MEIRLOP scans all input genomic regions for the presence of TF motifs. Then, it performs a dimensionality reduction using PCA on the k-mer composition of the input sequences and considers the loadings of the principal components that account for 99% of the variance. MEIRLOP then fits logistic regression models using the presence of the motif as a response variable and both the PC loadings describing the k-mer sequence composition and the input signal as predictor variables. For our benchmark, we used the log2 fold changes between the perturbed TF *vs.* control as input signal. While the resulting coefficients of the PC loadings describe associations between the presence of the motif to sequence composition (e.g., GC content), the coefficients of the input signal describe the association between the strength of the input signal and the presence of a motif. Wald test statistics on the model’s coefficients of the logistic regression are used to calculate p-values. Ranking of TFs is done based on the p-value and coefficients of the input signal in the logistic regression model.

Reference: *DeLos Santos et al.* MEIRLOP: improving score-based motif enrichment by incorporating sequence bias covariates. **BMC Bioinformatics**, 2020; 21 (410).

Software link: <https://github.com/npdeloss/meirlop>

7) MonaLisa

MonaLisa uses a randomized lasso stability selection regression approach. In monaLisa’s regression, the input signals are used as response variables. As input signal, we use in our benchmark uses the log2 fold change between the perturbed TF and the corresponding control. In monaLisa, the model matrix of predictor variables contains the number of TF motifs of each transcription factor (columns, predictors) in each genomic region (rows). MonaLisa incorporates two additional predictors in the model matrix with summaries of the CG and CpG composition of each genomic region. Thus, the randomized lasso stability selection model determines whether the presence of TF motifs can explain the input signal, accounting for sequence composition biases. Since the regression approach involves fitting for many random subsets of the data, monaLisa returns probabilities of motif selection and AUC values, which are used to rank TF motifs.

Reference: *Machlab et al.* monaLisa: an R/Bioconductor package for identifying regulatory motifs. **Bioinformatics**, 2022 March; 38 (9), 2624-2625.

Software link: <https://bioconductor.org/packages/release/bioc/html/monaLisa.html>

8) RcisTarget

RCisTarget uses a reference database of target genomic regions where TF motif have been extensively characterized. The characterization of TF motifs in these target genomic regions are constructed by considering motif clustering along the genome and their evolutionary conservation. Each target region in the database has a pre-computed ranked scored for each TF motif. RcisTarget inputs both a foreground set of genomic regions (differential peaks between perturbed TF and control, in our benchmark) and an optional background set of genomic regions. The program overlaps these input and background regions with the target regions in the database. Using the pre-computed ranking scores, it estimates an Area Under the cumulative Recovery Curve (AUC) for each TF motif using the foreground and background sequences. Afterwards, it standardizes the distribution of TF enrichment scores by subtracting the mean AUC value and dividing by the standard deviation of the AUC values. These NES values are used to rank the TFs.

Reference: *Aibar et al.* SCENIC: Single-cell regulatory network inference and clustering. **Nature Methods**, 2017; 14, 1083-1086.

Software link: <https://bioconductor.org/packages/release/bioc/html/RcisTarget.html>

9) TFEA

TFEA starts by defining a set of regions of interest (ROI). These are defined from the input regions (in our case, peaks in the foreground and the background conditions) by generating a probability distribution based on the read coverage and calculating joint probability distributions. ROIs positions are defined by the probability density function of the joint distributions. Using the regions of interest, a differential analysis test comparing the perturbed TF and the control is done to calculate a fold change and p-value. The log10 of the p-value, multiplied by the sign of the fold change is used as a rank statistic. For each TF motif, the rank statistic is weighted, with an exponential decay, as a function of the distance of the motif to the centre of the ROI. Then, an e-score is calculated by considering the area between the curve of cumulative sum of weighted ranks and a uniform background, scaled by the number of motif instances. To calculate a p-value of the motif enrichment, a null distribution of e-scores is estimated by reshuffling the weighted ranks for many times.

Reference: *Rubin et al.* Transcription factor enrichment analysis (TFEA) quantifies the activity of multiple transcription factors from a single experiment. **Commun Biol,** 2021; (4) 661

Software link: <https://github.com/Dowell-Lab/TFEA>

**Abbreviations:**

All tools were run in their default mode. For RcisTarget and HOMER, we also tested the options to provide a user-defined background. We also tested HOMER using as input the *Lambert* collection of motifs.

RcisTarget: default parameters using merged peaks from perturbed TF and control conditions.

RcisTarget + bg: default parameters using a user-provided background.

HOMER: default parameters.

HOMER + bg: default parameters with user-provided background.

HOMER + Lambert: default parameters with user-provided motif library.

HOMER + bg + Lambert: default parameters with user-provided motif library and user-provided background.

**Parameter optimization – Selected parameters and values**

**BART**

1. --nonorm

Whether or not do the standardization for each TF by all of its Wilcoxon statistic scores in our compendium. If set, BART will not do the normalization. Default = FALSE.

Tested values = [TRUE, FALSE].

**CRCMapper**

1. -l EXTENSION, --extension-length EXTENSION

Enter the length to extend subpeak regions for motif finding. Default = 100

Tested values = [50, 100, 250, 500]

2. -N NUMBER, --number NUMBER

Enter the number of non overlapping motifs in a region required to assign a binding

event. Default = 1

Tested values = [1, 2, 5, 10]

**HOMER**

1. Sequence normalization options:

-gc (use GC% for sequence content normalization, now the default)

-cpg (use CpG% instead of GC% for sequence content normalization)

-noweight (no CG correction)

Default = -gc

Tested values = [-gc, -cpg, -noweight]

2. -h (use hypergeometric for p-values, binomial is default)

Default = binomial

Tested values = [binomial, hypergeometric]

3. -nlen <#> (length of lower-order oligos to normalize in background, default: -nlen 3)
   Default = 3

Tested values = [0, 1, 2, 3]

**LOLA**

1. minOverlap

Minimum bases required to count an overlap. Default = 1

Tested values = [1, 2, 25, 50, 100]

2. redefineUserSets

run redefineUserSets() on your userSets? Default = False

Tested values = [True, False]

**MEIRLOP**

1. --kmer max_k

Set length of kmers to consider during regression. Principal components based on frequency of kmers will be used as a covariates in logistic regression. Set to 0 to disable. Default = 2

Tested values = [0, 1, 2, 3]

2. –length

Set this flag to incorporate sequence length as a covariate in logistic regression. Multiple covariates will be reduced to principal components.

Default = False

Tested values = [True, False]

3. --gc

Set this flag to incorporate GC content as a covariate in logistic regression. Recommend setting --kmer to 0 if using --gc. Multiple covariates will be reduced to principal components. Default = False

Tested values = [--kmer 0 –gc]

**MonaLisa**

1. Weakness

Value between 0 and 1. It affects how strict the method will be in selecting predictors. The closer it is to 0, the more stringent the selection. A weakness value of 1 is identical to performing lasso stability selection (not the randomized version). Default = 0.8

Tested values = [0.25, 0.5, 0.8, 1]

2. PFER

Integer representing the absolute number of false positives that we allow for in the final list of selected variables. For details see Meinshausen and Bühlmann (2010). Default = 2

Tested values = [1, 2, 5, 10]

**RCisTarget**

1. aucMaxRank

Threshold to calculate the AUC. In a simplified way, the AUC value represents the fraction of genes -within the top X genes in the ranking- that are included in the signature. The parameter 'aucThresholdPERC' allows to modify the percentage of genes (of the top of the ranking) that is used to perform this computation. By default it is set to 5% of the total number of genes in the rankings. Common values range from 1 to 10%. Default = 0.005

Tested values = [0.0025, 0.005, 0.01, 0.05, 0.1]

**TFEA**

1. --enrichment {auc,auc_bgcorrect}

Method for calculating enrichment. Default: auc

Tested values = [auc, auc_bgcorrected]

2. --largewindow LARGEWINDOW

The size (bp) of a large window around input regions that captures background. Default: 1500

Tested values = [1500, 3000, 6000]

3. --smallwindow SMALLWINDOW

The size (bp) of a small window arount input regions that captures signal. Default: 150

Tested values = [150, 300, 600]
